# Beta regression improves the detection of differential DNA methylation for epigenetic epidemiology

**DOI:** 10.1101/054643

**Authors:** Timothy J. Triche, Peter W. Laird, Kimberly D. Siegmund

**Author notes:** Present address: Center for Epigenetics, Van Andel Research Institute, Grand Rapids, MI 49503 USA. Corresponding author Email addresses: TJT PWL KDS.

## Abstract

**Background:** DNA methylation is the most readily assayed epigenetic mark, possessing confirmed relationships with gene expression, imprinting, and chromatin accessibility.Given the increasingly widespread use of DNA methylation microarrays in population-scale epidemiological applications, we sought to determine which methods provided the greatest statistical power to reproducibly detect differences in DNA methylation across various conditions,using publicly available data sets on tissue type and aging.

**Results:** Beta regression, as proposed originally by Ferrari and Cribari-Neto, yielded more validated hits in each of our comparisons than any other method under consideration, both in a regression setting and in comparisons to two-group tests such as the Wilcoxon-Mann-Whitney, Student t, and Welch t tests.In large cohorts of whole blood samples, we corrected for compositional differences and batch effects, and found that marginal likelihood ratio tests from beta regression models uniformly dominate popular alternatives based on linear models.The superior sensitivity and specificity exhibited by beta regression in epidemiologically relevant cohort sizes corresponded to approximately a 2% increase in sensitivity at the same specificity when compared to linear models fitted on raw beta values (proportion of signal intensity due to the methylated allele), M-values, or rank quantile normalized values.

**Conclusions:** Investigators should consider beta regression to maximize statistical power in studies of DNA methylation using microarrays.At epidemiologically relevant sample sizes, with typical quality control procedures (compositional and batch effect correction), cross-cohort agreement uniformly favors beta regression over popular alternatives.

## Introduction

Epigenetic information - modifications to chromatin and its constituent DNA, as opposed to genetic information encoded in the DNA sequence itself - is now widely recognized as playing an important role in development, degeneration, and disease. In vertebrate genomes the most readily accessible epigenetic mark is cytosine methylation at CG dinucleotides, associated with transcriptional repression when found in promoter regions. In population-scale studies of multifactorial outcomes, improved sensitivity to detect true associations between DNA methylation changes and exposures or outcomes is of great interest. We therefore sought to explore the trade-offs involved in faithfully modelling the proportional measurements via maximum likelihood beta regression [<u>1</u>], vs. least-squares modelling after logit transformation [<u>2</u>], or a N(0,1)-quantile normalization [<u>3</u>], or no transformation.

Typical approaches to modelling DNA methylation at individual cytosines include ignoring the heteroskedasticity of proportional (0-1) measurements; N(0,1)-quantile normalizing the measurements [<u>3</u>]; logit-transforming the measurements and working on a fold-change scale [<u>2</u>], or discretizing proportions into “states” (for example, regions of unmethylated, variably methylated, and highly methylated cytosines) which are presented as surrogates for biologically relevant changes [<u>4</u>]. The latter discretization typically assumes extensive coverage (as with whole-genome bisulfite sequencing), making strong assumptions regarding local correlation on the genome. By contrast, we focus on regression approaches useful for sensitive modelling of associations at individual targeted cytosines, as a function of exposures, which could be continuous (e.g. age) and typically require sample sizes that would be cost-prohibitive for whole-genome bisulfite sequencing. We evaluated Beta regression and common approaches to linear models, with or without transformation, for various sample sizes and designs.

Linear models are quite flexible and can be solved with relative ease even for highly underdetermined datamatrices (e.g. via penalized least-squares methods). Similarly, any generalized linear model can be iteratively solved to find constrained maximum likelihood estimates of predictor effects on a response, with certain distributional assumptions. Beta regression, as proposed by [<u>5</u>, <u>6</u>], provides a regression framework that will be familiar to practitioners comfortable with generalized linear models. However, the model allows for the coupled mean and variance terms to both depend on covariates. It is not feasible, with a single uniformly applied transformation across responses, to simultaneously normalize the proportional measurements and completely decouple its variance from its mean (although transformations which render the data approximately homoskedastic, e.g. [<u>2</u>], have proven useful). Thus, directly modelling both terms as a function of the data is an attractive approach.

In addition, changes in the variability of DNA methylation are also recognized as predictors of interest for many diseases [<u>7</u>] and more recently, as nearly inevitable side effects of aging [<u>8</u>]. A dispersion (variability) parameter is integral to the beta regression model, permitting either or both of the mean and dispersion to depend upon covariates. Moreover, one of the most popular biospecimens for population-scale epigenomics work - whole blood - is in fact a complicated mixture of myeloid, lymphoid, and dendritic cells [<u>9</u>], each with characteristic DNA methylation marks [4]. While cell sorting is a useful method to reduce this ambiguity, it is not always feasible to separate fractions (e.g.[<u>10</u>]). Greater sensitivity is thus useful for detecting phenotypic associations of interest, which may only be present in a fraction of cell types. At the same time, regression analysis allows us to model the influence of other factors, such as fraction of cell type (e.g. [<u>11</u>, <u>12</u>]), which may influence the estimated associations. Therefore, we employed several publicly available data sets to explore various approaches to modelling variation in DNA methylation.

First, we evaluated Beta regression in the context of dichotomous exposure. We used the myeloid/lymphoid classification of sorted blood cell fractions to evaluate common two-group tests (Student's t test, the Welch t test, Mann-Whitney-Wilcoxon) on variously transformed or untransformed data, and to compare the sensitivity of these to that of likelihood ratio tests based on the Beta regression model, with either a shared estimate of variability (the “Beta-T” test) or a fitted per-group estimate of variability (the “Beta-Welch” test).

Second, we analyzed two large studies associating aging, as a continuous exposure, with DNA methylation levels in whole blood [<u>13</u>][<u>14</u>]. These large studies allowed us to use one (HuGeF) as a discovery set and the other (Hannum) as a test set when gauging sensitivity of different methods. As we know blood to be a complex mixture of different cell types varying with age, we used the compositional estimation procedure from Houseman [<u>15</u>] in addition to factors for batch, plate, gender, and ethnicity (where the latter was available) to reasonably approximate the steps which would be expected in an epidemiologically relevant cohort study. Sensitivity and specificity was estimated as the proportion of “true positives” (replicated hits) at a given specificity (putative error level in the discovery cohort) and plotted in Fig.1A

## Methods

### Models for differential DNA methylation

The ‘betareg’ R package (version 3.0.4), as described in [<u>1</u>], was employed for the Beta regression fits. Instead of the usual formulation, Beta(a,b), with a, b>0, mean μ=a/(a+b) and variance σ^2^=μ(1- μ)/( a+b+1) as a function of the mean, the reparameterized distribution Beta(μ,φ) is used, where φ=a+b is a precision parameter, independent of the mean. For small φ, the variance is large, and for large φ,the variance is small. Briefly, given link functions g_1_(μ) and g_2_(φ), we model the mean μ and precision φ as a function of predictor matrices X and Z, respectively, to find maximum likelihood estimates of the coefficients β and γ. Assume the Beta value for locus *j* from individual *i* is y_ij_ ~ Beta(μj,φj). For simplicity, we drop the index *j.* Then,

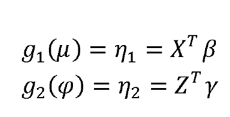

with the links for g_1_(μ) and g_2_(φ) being logit(μ) and log(φ), respectively. The log-likelihood is then expressed for responses of N subjects as

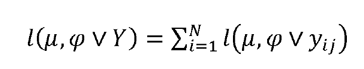

such that each individual observation contributes to the loglikelihood the quantity

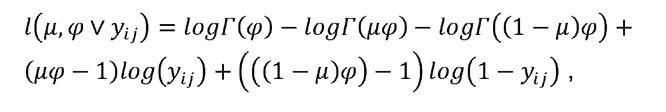
where Γ is Euler's Gamma function (Γ(n)=(n-1)!).

We implemented two changes that we found necessary to improve speed and stability. First, we employed the “squeezing” technique suggested by [<u>6</u>] to eliminate 0 and 1 values without altering the rank order of observations. To squeeze the data we shifted the values around zero and multiplied by 0.99 (=(Beta - 0.5)*0.99+0.5). Second, we used an approach similar to that of ‘limma’ [<u>18</u>], with predefined design matrices for the null and alternative hypotheses, for model fitting. Fitting a 2-3 variable model to each CpG site on the HumanMethylation450 array took under 6 hours on a machine with 6 processors and 48GB of RAM, when a likelihoodratio test was conducted at each locus.

For linear regression, the ‘limma’ package was employed, using the marginal t-test statistics for each coefficient of interest. Models were fit using untransformed and transformed Beta values. Untransformed Beta values, the ratio of intensities M/(M+U), where M measures methylation and U lack of methylation intensities, were labelled ‘(B)’ or ‘betas’.We also considered M-values, log2(M/U), labelled ‘(M)’, owing to their purportedly superior sensitivity, as described in [<u>2</u>]. Lastly, we evaluated the normalizing transformation described in [<u>3</u>], whereby the observed quantiles at a given locus are replaced with their corresponding values for a standard Normal distribution (i.e., N(0,1)) labelled ‘(NQ)’.

### Error rate control

We used independent data sets to validate loci associated with age, allowing us to compare sensitivity of the different analysis approaches. For the blood lineage comparisons, a validation set was not available. For this data set, we permuted the samples, blocked by subject, showing control of the type I error through permutation.

### Data sets

All analyses in this paper were applied to publicly available data. When available (in fractionated normal blood cells and the HuGeF-EPIC Italy cohort), raw IDAT files were downloaded and processed using the ‘methylumi’ package to correct background fluorescence and equalize within-array dye-bias[<u>19</u>].Where raw data was not available, internal control probes were used by GenomeStudio to equalize dye bias and correct background fluorescence. Probes were mapped to genomic coordinates using manufacturer annotations verified in the ‘FDb.InfiniumMethylation.hg19’ package, masking common SNPs, repeat regions, and sex chromosome probes. This mask is also used for the Cancer Genome Atlas project HumanMethylation450 data, and excludes probes with common SNPs at, or up to 15 bases 3’ to, the interrogated locus, as well as probes with at least 15 bases proximal to the interrogated locus falling into an annotated repeat region of the genome. Regressions were fit separately per probe (excluding masked loci), and (as noted) p-values were adjusted using Benjamini & Hochberg's procedure [<u>20</u>] to control the FDR at 0.05.

A data set of 48 samples of sorted cell fractions (24 myeloid/ 24 lymphoid samples) from 6 adult male subjects was used to compare methods for testing differential DNA methylation in two-group designs (GEO series GSE35069) [<u>12</u>]. Reinius et al. (2012)[<u>12</u>] showed that cell type-specific differences dominated individual-specific differences in these samples, so we modelled DNA methylation as a function solely of cell type (myeloid or lymphoid). IDAT files were downloaded from (<u>http://publications.scilifelab.se/kere_j/methylation.</u>) and processed as described above.

Two data sets, one from the HuGeF-EPIC Italy consortium (unpublished, GEO GSE51032, N=845) and one from Hannum (2013, GEO series GSE40279, N=656), were used to test for association between CpG methylation and age, a quantitative variable. We fitted models including age, gender, plate, batch, ethnicity (in Hannum only, not available for HuGeF) and estimated cell counts (monocytes, granulocytes, B-cells, CD4+ T-cells, CD8+ T-cells, CD56+ NK cells) to each dataset, using the HuGeF-EPIC Italy dataset for discovery and Hannum for validation.

## Results

Table 1 presents the number of loci discovered to have differential DNA methylation in myeloid and lymphoid cell fractions, in a data set with 48 samples of sorted blood cells from 6 healthy males. The “Beta-T” and “Beta-Welch” tests were most sensitive among the methods compared, discovering 194,767 (51%) and 204,189 (54%) out of 381,552 loci. However, in these data, the marginal gain over the number of loci discovered by any other approach was modest. A large number of loci were identified by all approaches (177,446 loci, 91% of the loci discovered by “Beta-T” and 87% of those found using “Beta-Welch”). Permutations confirmed that the error rate was adequately controlled by all methods. Although there was variation in empirical error rate between the two permutations, this is likely a function of the small sample size (6 subjects). The rank order of the methods was not altered when we computed either Benjamini-Hochberg FDR-adjusted p-values or q-values for multiple comparisons correction (data not shown).

**Table 1.**
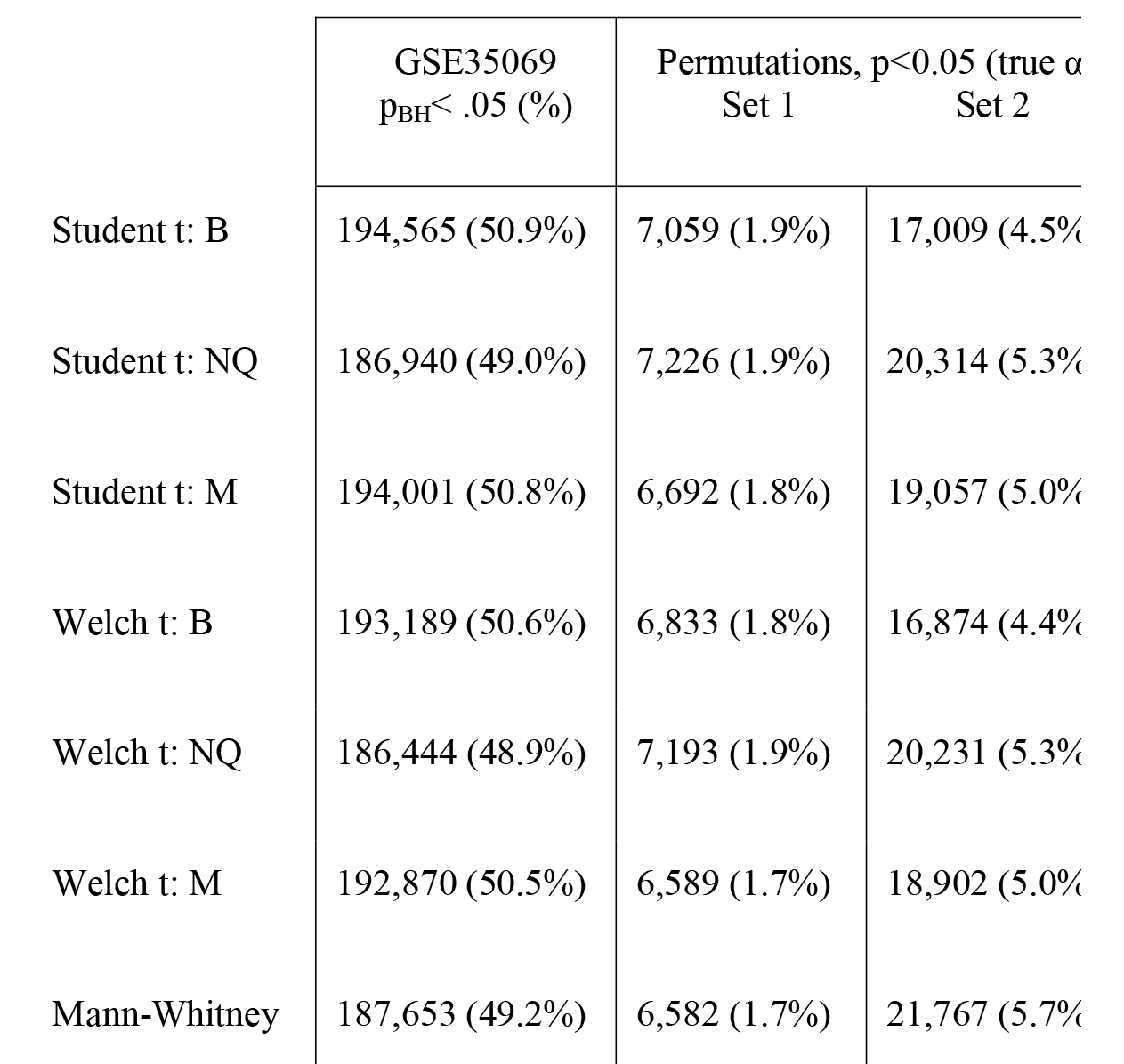
Autosomal loci associated with lineage in blood

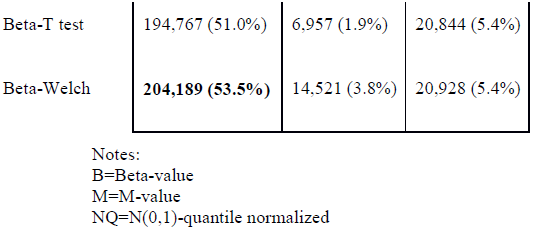

> Set 1 and Set 2 are two different permutation data sets, obtained by swapping lymphoid and myeloid samples for three of six subjects, maintaining the original balance of ages (verified by personal correspondence with Dr. Kere) between the donors.

Figure 1 presents the relative sensitivity and specificity (judged by cross-cohort replication at a given p-value cutoff in the discovery cohort) for the various methods under consideration, after correcting for cellular composition (the same figure constructed without correcting for cellular composition is presented as supplementary figure 1). We note that beta regression uniformly outperforms all competing methods herein.

**Figure 1:**
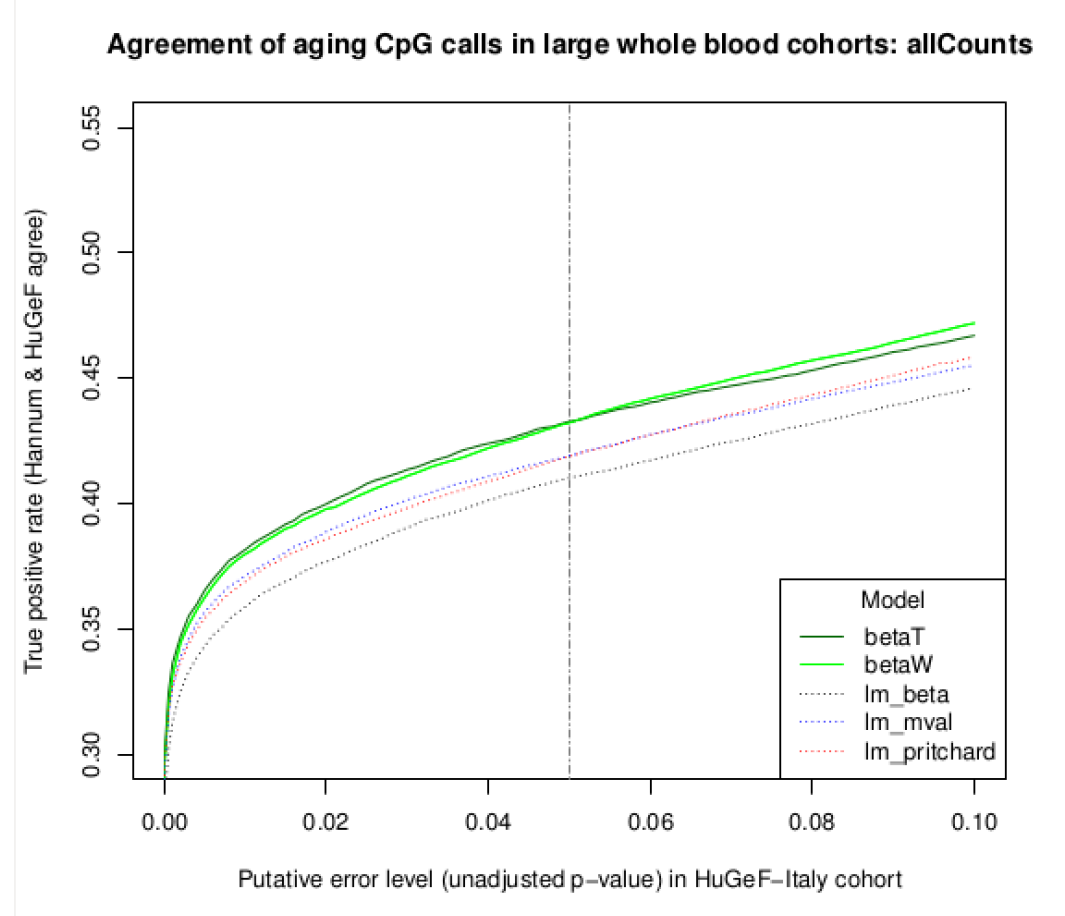
Replication (“true positive”) versus putative error rate (“false positive”) for methods compared in Results.

## Discussion

As a relatively young field, genome-scale investigation of DNA methylation (and indeed epigenetic marks in general) has spawned multiple approaches to assess the significance of focal differences in DNA methylation (1bp-1kbp), as well as broader concordant DNA methylation patterns, often associated with chromatin accessibility states. Both focal and broad changes may be associated with a phenotype of interest, and both are strongly associated with tissue-specific changes involved in development and disease.

Our results suggest that, for data from Illumina Infinium methylation chips, Beta regression offers the greatest sensitivity to detect these changes in DNA methylation, and for the analysis of age-related changes, this result carried through to validation in independent datasets. However, whole blood is known to be a mixture of cell types and the fractions of those cell types vary over age [<u>16</u>]. Therefore it is expected that a number of the validated loci may not have differences in DNA methylation with age within individual cell fractions, but rather it is the changing composition of cells types with age that allow cell-type specific marks to result in different average DNA methylation levels with aging. Thus caution is needed when interpreting results from complex cell mixtures.

If, as reported for whole blood, cell type-specific aging is faster in one or more cell fractions than the majority of cells [<u>15</u>], one might expect the cell-type specific DNA methylation differences with age to be attenuated in the mixed cell population. We therefore corrected for the changing estimated cell counts at the single-sample level, and found that this decreased both the overall number of hits and the proportion of replicated hits across blood cohorts, suggesting that its proposed critical importance is indeed a valid concern in the analysis of studies of epigenetic epidemiology.

The data sets we selected for analysis had large sample sizes, as will be typical in an epidemiologic genome-wide DNA methylation study. In previous work, we conducted a simulation study to evaluate control of the false-discovery rate in small samples for the methods evaluated in this paper (see Supplemental file 1). These results indicated that very small (<25 samples per group) or unbalanced comparisons (splits more uneven than approximately 80%/20%) designs, could lead to loss of control of the error rate for Beta regression. This might be due to instability of the unconstrained maximum likelihood estimates in small samples. Various approaches to bias correction and reduction have been proposed [<u>1</u>], but these can increase computation times by at least an order of magnitude. In order to establish control of the error rate for the (unvalidated) two-group tests analyzed in this paper, having different genomic distribution and potentially different properties than studied in our simulations, we analyzed two permutations of the samples, randomly switching the group assignments for age-balanced sets of subjects. We recommend this type of verification for error-rate control when an independent data set is not available.

As always, the investigator must decide what the goal of the experiment or study should be, and what the most appropriate tools may be to accomplish the goal. Computational resources and time frames for completion can guide such choices, but ultimately the nature of a particular study should determine the method(s) of analysis. Exploratory analyses may be better served by the speed of linear methods, which allow for rapid iteration of models fitted to millions of data points.

By contrast, an additional day of computation may not be a significant consideration for a cohort study investigating changes in DNA methylation at specific loci in normal tissues as associated with exposures, where sensitivity might well be a central consideration, and all resulting findings subject to independent orthogonal validation. Such situations may favor sensitivity over speed of computation.

Similarly, when characterizing regional changes at a given significance threshold, either for a sequential area statistic or by modelling smoothed runs of multiple test statistics, additional sensitivity may help to unearth biologically interesting patterns that are difficult to discern otherwise. Future work will investigate an omnibus test statistic for both variable and fixed effects which appears to outperform competitors, and may yield more fruitful leads than individual tests of either type. Nonetheless, the sequential nature of regional differential methylation calls requires permutation to establish a robust FDR estimate, and this permutation step itself provides additional control of the putative error rate at the regional level. Therefore, we contend that greater sensitivity at the local (individual CpG) level is of interest even if broad, subtle effects are anticipated.

## Conclusions

Beta regression offers greater sensitivity for the discovery phase of genome-wide DNA methylation association studies, particularly in large sample sizes with potentially confounding per-group effects on dispersion. The Beta regression framework appears to draw its enhanced power from the natural representation of such effects, which is more challenging in a linear model due to the decoupling of the variance from the mean. The ability of the Beta regression framework to simultaneously model changes in the mean and dispersion grants a significant advantage when modelling factors (such as aging) that appear to influence both quantities (and indeed confounding factors such as cellular composition). Investigators dissecting complex epigenetic associations with continuous phenotypes should include beta regression in their tools for discovery, balancing computational convenience and conservative error rates of linear modelling approaches against the need for sensitivity and flexibility in modelling dispersion.

## Acknowledgements

Research reported in this publication was supported by the National Institutes of Health: National Human Genome Research Institute Award Number R01HG006705 (KS) and National Cancer Institute Award Numbers R01 CA097346 (KS) and R01 - CA170550 (PWL). The content is solely the responsibility of the authors and does not represent the official views of the National Institutes of Health. The authors would like to acknowledge Juha Kere, for IDAT files of his data; and the USC High Performance Computing Center, for computational resources.

## Authors' contributions

TJT designed and conducted the analyses, and wrote the manuscript. KDS conceived of the experiment and performed a pilot study in a separate dataset. PWL supervised the analyses and offered insights regarding the biological relevance of the results.

